# Reduced dopamine signaling impacts pyramidal neuron excitability in mouse motor cortex

**DOI:** 10.1101/2020.02.12.946301

**Authors:** Olivia K. Swanson, Rosa Semaan, Arianna Maffei

**Author notes:** **Contributions.** O.K. Swanson designed and performed experiments, analyzed data; R. Semaan performed immunohistochemistry and stereological quantification of lesions; A. Maffei designed and supervised the experiments and analysis. All authors were involved in writing and commenting the manuscript. **Send correspondence to:** Arianna Maffei, Dept. of Neurobiology and Behavior, Life Science Building Rm 548, SUNY – Stony Brook, Stony Brook, NY 11794-5230.

## Abstract

Dopaminergic modulation is essential for the control of voluntary movement, however the role of dopamine in regulating the neural excitability of the primary motor cortex (M1) is not well understood. Here, we investigated two modes by which dopamine influences the input/output function of M1 neurons. To test the direct regulation of M1 neurons by dopamine, we performed whole-cell recordings of excitatory neurons and measured excitability before and after local, acute dopamine receptor blockade. We then determined if chronic depletion of dopaminergic input to the entire motor circuit, through a mouse model of Parkinson’s Disease, was sufficient to shift M1 neuron excitability. We show that D1 and D2 receptor (D1R, D2R) antagonism altered subthreshold and suprathreshold properties of M1 pyramidal neurons in a layer-specific fashion. The effects of D1R antagonism were primarily driven by changes to intrinsic properties, while the excitability shifts following D2R antagonism relied on synaptic transmission.

In contrast, chronic depletion of dopamine to the motor circuit with 6-hydroxydopamine (6OHDA) induced layer-specific synaptic transmission-dependent shifts in M1 neuron excitability that only partially overlapped with the effects of acute D1R antagonism. These results suggest that while acute and chronic changes in dopamine modulate the input/output function of M1 neurons, the mechanisms engaged are distinct depending on the duration and location of the manipulation. Our study highlights dopamine’s broad influence on M1 excitability by demonstrating the consequences of local and global dopamine depletion on neuronal input/output function.

**Significance statement:** Dopaminergic signaling is crucial for the control of voluntary movement, and loss of dopaminergic transmission in the motor circuit is thought to underlie motor symptoms in those with Parkinson’s Disease (PD). Studies in animal models of PD highlight changes in M1 activity following dopamine depletion, however the mechanisms underlying this phenomenon remain poorly understood. Here we show that diminished dopamine signaling significantly alters the excitability and input/output function of M1 pyramidal neurons. The effects differed depending on the mode and location – local versus across the motor pathway – of the dopamine manipulation. Our results demonstrate how loss of dopamine can engage complex mechanisms to alter M1 neurons activity.

## Introduction

Primary motor cortex (M1) exerts powerful control over movement execution through its central location in the motor circuit. It receives inputs from other cortices and the thalamus, the latter relaying converging signals from the basal ganglia and cerebellum, (Mink, 1996; Bosch-Bouju et al., 2013; Hooks et al., 2013) and makes direct connections to descending motor tracts (Lemon, 1993). Pyramidal neurons in each layer of M1 show distinct projection patterns, with neurons in the superficial layers largely innervating deep layers of M1, other cortices, or striatum; neurons in deep layers primarily project back to the thalamus, striatum, or to the corticospinal tract (Oswald et al., 2013). Integral to the transition from input to motor output is neural excitability, which influences the magnitude of a neuron’s response to incoming activity. Excitability can be modulated by motor learning and synaptic plasticity, as well as changes in overall synaptic drive (Paz et al., 2009; Kida et al., 2016). Neuromodulators further contribute to regulating excitability and input/output functions of neurons, and the most crucial neuromodulator for the central control of movement is dopamine.

Dopamine’s influence on M1 excitability remains poorly understood (Vitrac and Benoit-Marand, 2017). Anatomical studies report that dopaminergic neurons extend projections to M1 (Fallon and Moore, 1978; Williams and Goldman-Rakic, 1998; Hosp et al., 2011), and dopaminergic activity modulates M1 neurons firing rates (Vitrac et al., 2014). Direct dopaminergic projections are most dense in the deep layers of rodent M1 (Nomura et al., 2014), and D1-like and D2-like receptors (D1R, D2R) are expressed along the entire depth of the cortical column with laminar-specific density (Ariano and Sibley, 1994; Khan et al., 1998; Lemberger et al., 2007), suggesting possible laminar differences in the effects of local dopamine modulation.

However, the largest dopaminergic input to the motor circuit projects from the substantia nigra pars compacta (SNc) to the basal ganglia (Beckstead et al., 1979), where it signals through D1R and D2R with opposing effects on neural excitability (Surmeier et al., 2007; Azdad et al., 2009; Planert et al., 2013). This, along with its control of synaptic plasticity and transmission, make dopamine a powerful regulator of basal ganglia output (Bagetta et al., 2010) with the ability to affect activity across the motor circuit.

Diminished dopaminergic input to the motor circuit can profoundly impair movement, as observed in Parkinson’s Disease (PD). While it is clear that depleted dopamine leads to shifts in excitability and synaptic transmission (Blandini et al., 2000; Jankovic, 2008; Grieb et al., 2013; Calabresi and Di Filippo, 2015), this evidence is largely restricted to studies of the basal ganglia. It is unclear if altered basal ganglia activity following nigral dopamine depletion impacts the input/output function of M1 neurons. The classic model of PD postulates that loss of dopamine in the basal ganglia increases the inhibitory output of these nuclei and leads to decreased activation of M1 (Albin et al., 1989; McGregor and Nelson, 2019), which could consequently alter excitability. Furthermore, patients with PD exhibit reduced dopaminergic axon density directly in M1, and functional studies of patients as well as animal models of the disease show altered M1 neural activity, suggesting that depleted dopamine within M1 could also play a role in shifting M1 neural excitability (Gaspar et al., 1991; Lefaucheur, 2005; Lindenbach and Bishop, 2013).

Here, we examined the effect of acute and chronic loss of dopamine signaling on the input/output function of M1 pyramidal neurons using patch clamp recordings in acute brain slices. First, we tested how acute blockade of D1R or D2R affects neuronal excitability in superficial and deep layers of M1. We then asked whether chronic loss of dopaminergic signaling either in the midbrain or locally within M1 impacts M1 neural excitability. Our results show laminar-specific effects on M1 input/output function through multiple mechanisms. Acute antagonism of D1R impacted excitability through modulation of intrinsic properties, while the effect of D2R blockade depended on synaptic transmission. Chronic depletion of dopaminergic neurons in the midbrain was also sufficient to engage synapse-dependent modulation of M1 neurons input/output function, although the effects only partially overlapped with those observed following acute blockade. These data show that loss of dopamine impacts M1 neural excitability and highlights the complex mechanisms that can be engaged depending on laminar location within M1 and specific manipulation of dopamine signaling.

### Experimental procedures

Surgical and experimental procedures followed the guidelines of the National Institute of Health and were approved by the institution’s Animal Care and Use Committee. C57BL/6 mice of both sexes were used in the following experiments. Number of animals and number of cells in each experimental group were shown as *N* and *n*, respectively, in the figure legends.

#### 6OHDA injection

Chronic dopamine depletion was achieved by injection of 6-hydroxydopamine hydrochloride (6OHDA) either in the SNc or in M1. Prior to surgery, animals received an IP injection of desipramine (1.25mg/mL, 20mL/kg) to protect noradrenergic and serotonergic afferents from taking up 6OHDA. C57BL/6 mice of both sexes (P35-P45) were anesthetized (100mg/kg ketamine and 10mg/kg xylazine) and a craniotomy was made over the injection site of interest. The 6OHDA solution (15mg/mL in 0.02% ascorbate) was prepared fresh at the time of injection. Mice unilaterally injected within M1 received 3μg 6OHDA, or vehicle, via two 100nL injections (Bregma+1.2/0.8, midline-1.1, surface-0.8). Animals unilaterally injected in the SNc received 7.5μg 6OHDA, or vehicle, via two 250nL injections (Bregma-3.1/2.8, midline-1.2, surface-3.93). A pressure injection system (Nanoinject, Drummond) was used for these procedures. Following surgery, animals were monitored daily for food and water intake, and administered fluids or softened food when needed.

#### Cylinder motor task

Prior to slice preparation, all 6OHDA or vehicle-injected animals were assessed for motor impairment using the cylinder motor task (Iancu et al., 2005). The animal was placed in a clear acrylic cylinder and allowed to freely explore for 10min, while being filmed with a camera positioned on top of the cylinder. Mirrors were positioned around the cylinder to facilitate visualization of forelimb use and post hoc analysis (see Fig. 4). Weight-bearing forelimb wall touches were counted over a 3-minute period, or a minimum of 20 touches, by an experimenter who was blind to the surgical procedures. Use of forelimb was quantified as a ratio of wall touches by the forelimb contralateral to the injection (vehicle or 6OHDA) over total wall touches.

#### Slice electrophysiology

Animals at P50-P70, or 2-3 weeks after surgery for vehicle and 6OHDA injected groups, were deeply anesthetized with isoflurane using the bell jar method and rapidly decapitated. Following dissection of the brain, acute 300μm slices containing forelimb M1 (Tennant et al., 2011) were prepared using a vibrating blade microtome (Leica VT1000S). Tissue was sectioned in ice cold oxygenated ACSF, recovered for 30min in 37°C ACSF, then allowed to stabilize at room temperature for at least 40min prior to recording. Whole-cell patch clamp of visually-identified excitatory neurons was performed at room temperature using pulled borosilicate glass pipettes with a resistance of 3-4MΩ). Dynamic input resistance and frequency-current measurements were obtained in current clamp by injecting 700ms current steps of increasing intensity (−100 to 450pA at 50pA increments). Action potential threshold was measured on single action potentials at rheobase (rheobase was determined as the current step, in 2pA increments, that elicited a single action potential). Voltage dependence of the I_h_ current was measured in current clamp, as the amplitude of the voltage sag current induced by 700ms hyperpolarizing current steps from −200pA to −25pA, in 25pA increments. To block dopamine receptors, either a D1 (SCH23390) or a D2 (sulpiride) receptor antagonist was bath-applied for 15min following a 10min baseline. To assess the dependency of dopaminergic activity blockade on synaptic transmission, bath application of D1 or D2 receptor antagonists was repeated in the presence of fast synaptic transmission blockers (APV, DNQX and picrotoxin). Synaptic transmission blockers were circulated for 10min prior to the application of the dopamine receptor antagonists, and during this time spontaneous activity was monitored in voltage clamp to ensure that all synaptic events onto the recorded cell were abolished. Series resistance (Rs) was monitored throughout all experiments and data from cells with Rs>10% of input resistance, or changing >20% throughout the recording, were excluded from the analysis.

#### Immunohistochemistry

Recorded slices, along with the remaining brain tissue, were post-fixed in 4% paraformaldehyde (0.01M phosphate buffered saline (PBS), pH 7.4) for a minimum of one week. Remaining brain tissue containing injection sites was sectioned in the coronal plane at 50μM using a vibrating blade microtome (Leica VT1000S) and stored in PBS at 4°C. Recorded slices were rinsed in PBS and incubated for 30min at 45°C in an antigen retrieval solution (10mM sodium citrate, pH 8.5). Slices were again rinsed in PBS, incubated for 2h in 50mM glycine at room temperature (RT), then following an additional rinse, were pre-blocked for 3h at RT in PBS containing 5% bovine serum albumin (BSA, Sigma-Aldrich), 5% normal goat serum (NGS, Vector), and 1% Triton-X (Tx, VWR). Slices were then incubated overnight at 4°C in an antibody stock solution containing PBS, 1% BSA, 1% NGS, 0.1% Tx, and the following antibodies: Streptavidin Alexa Fluor 568 (1:2000, Invitrogen S11226) and mouse anti-GAD67 (1:500, EMD Millipore MAB5406). Slices were then rinsed and incubated for 6h at RT in the same antibody stock solution, containing goat anti-mouse Alexa Fluor 647 (1:500, Invitrogen A-21235), and were counterstained with Hoechst33342 (1:5000, Invitrogen H3570). Following an additional rinse in 0.1M phosphate buffer (PB), slices were mounted on gelatin-coated slides, and coverslipped with fluorescent mounting medium (Fluoromount-G, ThermoFisher).

To assess dopamine neuron or bouton loss, free-floating sections containing vehicle or 6OHDA injection sites were rinsed, then incubated in antigen retrieval solution and glycine as described above. Sections were incubated in 0.3% hydrogen peroxide (Fisher Scientific) for 30min, rinsed, and endogenous avidin and biotin were blocked (Avidin/Biotin blocking Kit, Vector). Following an additional rinse, sections were pre-blocked in the previously described solution with 0.2% Tx for 1h. Sections were then incubated overnight at 4°C in the antibody stock solution with 0.1% Tx and rabbit anti-tyrosine hydroxylase (1:1000, Abcam ab112), rinsed, and incubated for 3h at 4°C in the antibody solution containing biotinylated goat anti-rabbit (1:200, Vector BA-1000). Sections were rinsed and incubated in avidin-biotin horseradish peroxidase (Vectastain Elite ABC kit, Vector) for 1hr at RT, rinsed, and developed for 60s in 3,3’-diaminobenzidine (DAB Peroxidase Substrate Kit, Vector). At the end of this process, sections were rinsed in PB, mounted on gelatin-coated slides, and air-dried. Slides were then dehydrated in a series of alcohols (70%, 95%, 100%), cleared in xlyenes, and coverslipped with Entellan mounting medium. Imaging of fluorescently labeled sections was performed on a laser-scanning confocal microscope (Olympus), and brightfield images were obtained using a widefield microscope (Olympus).

For determination of cortical layers, a subset of animals was transcardially perfused first with PBS followed by 4% paraformaldehyde. Brains were dissected and post-fixed for 24 hours in 4% paraformaldehyde then sectioned as described above. 50μm sections containing M1 were stained with the same methods as above for the expression of the cytoarchitectural marker SMI-32 (mouse anti-SMI32, 1:2000, Biolegend 801701; goat anti-mouse Alexa Fluor 488, 1:500, Thermo Fisher A-11001), and counterstained with a pan-cellular nuclear stain (Hoechst33342, 1:5000, Invitrogen H3570) and neuronal-targeting fluorescent Nissl stain (Neurotrace530/615, 1:200 Thermo Fisher N21482).

#### Stereological analysis

The effect of vehicle or 6OHDA on dopaminergic neurons in the SNc and ventral tegmental area (VTA), and of putative dopaminergic boutons in M1 were assessed using unbiased stereological methods with the Stereoinvestigator system (Microbrightfield) (Gundersen, 1986; Grieb et al., 2013). These assessments were performed by an experimentalist who was blind both to surgical procedures and electrophysiological results. Sections containing injection sites were processed as described for tyrosine hydroxylase (TH), which labels dopaminergic neurons in the midbrain. Contours of the SNc, VTA, and layers in M1 were traced based on cytoarchitectural bounds from Nissl-stained adjacent sections, combined with chemoarchitectonic delineations as previously described (Fu et al., 2012). For counts of TH^+^ neurons in the SNc, 8 sections spaced 150μm apart were counted at 400x magnification, using a grid size of 150μm×150μm and a 100μm×100μm counting frame. The dissector height was set to 20μm with a guard zone of 2μm. Counts of TH^+^ neurons in the VTA were performed in the same manner, with 6 sections per animal. TH^+^ axon varicosities (putative boutons) were counted in layer 2/3 and layer 5, across 2 consecutive 50μm sections at 1000x magnification, using a grid size of 100μm×100μm and a 40μm×40μm counting frame. The dissector height was set to 20μm with a guard zone of 2μm. These sampling parameters were sufficient to yield population estimates with a coefficient of error (Gunderson CE) less than 10% for the un-lesioned hemispheres and were applied to all cases used in the study. Lesion severity was expressed as the estimated population of TH^+^ neurons in the injected hemisphere’s region of interest, relative to that of the contralateral hemisphere. Animals with lesion quantification falling beyond two standard deviations of the mean were excluded.

#### Data Analysis

Analysis of electrophysiological data was performed with custom-made procedures in Igor (Wavemetrics). Dynamic input resistance was computed as the slope of the current-voltage (IV) curve obtained from a series of hyperpolarizing current steps (−100pA to 0pA). Rheobase was determined by injecting current steps at 2pA increments until reaching the generation of a single action potential. Action potential threshold and rise time were both calculated at rheobase, in the following manner: First, the x-axis coordinate of the last zero crossing preceding the max of the second derivative of the trace was calculated. This x position was then applied to the original trace, and the threshold was determined to be the y-value at this point. For rise time, first the action potential amplitude was calculated as the difference in membrane potential between the peak of the action potential and threshold. The action potential rise time was calculated as the duration between the points of 10% and 90% amplitude on the rising phase of the action potential. Frequency-current (*fI*) curves were computed as the average frequency of action potentials for a given current step across all cells in each experimental group (50pA to 450pA). Voltage-dependence of I_h_ was measured as the difference in membrane potential between absolute minima of the membrane potential within the first 300ms of the current step and the average of the steady-state portion of the current step (the last 200ms) across hyperpolarizing steps.

#### Solutions

Artificial Cerebrospinal Fluid (ACSF) used in all electrophysiology experiments contained the following (in mM): 126 NaCl, 3 KCl, 25 NaHCO_3_, 1 NaHPO_4_, 2 MgSO_4_, 2 CaCl_2_, and 14 dextrose. Internal solution contained the following (in mM): 100 K-Glu, 10 K-HEPES, 4 Mg-ATP, 0.3 Na-GTP, 10 Na-phosphocreatine, and 0.4% biocytin, pH 7.35 titrated with KOH and adjusted to 295 mOsm with sucrose (E_rev_[Cl^−^] = −49.8mV). D1 and D2 receptor antagonists SCH23390 and (S)-(-)-Sulpiride (Tocris) were prepared in DMSO and diluted in ACSF to a final concentration of 10μM. Solutions containing these antagonists were kept in the dark and bath-applied during recording. Experiments performed in the presence of synaptic transmission blockers used ACSF containing the following (in μM): 20 DNQX, 50 AP5, 20 Picrotoxin. The maximum concentration of DMSO for any condition was 0.3% (synaptic blockers with D1 or D2 antagonist), however the transition from pre- to post-dopamine receptor antagonism did not exceed an increase of 0.1% DMSO.

#### Statistical Analyses

Statistical tests were performed in Microsoft Excel and the add-in statistical program XLSTAT. All data are presented as mean ± standard error of the mean (SEM) for the number of neurons (*n*) and the number of animals (*N*) indicated. All data were tested for normal distribution using the Shapiro-Wilk test. 2-tailed Student’s t-tests, paired or unpaired, were used to test for significance when appropriate (measurements of input resistance, threshold, rise time, cylinder motor assessment, stereological counts). The non-parametric Wilcoxon signed-rank test or Mann-Whitney U-test were used for data that did not follow a normal distribution (voltage dependence on I_h_, frequency-current curves). P-values ≤0.05 were considered significant.

## Results

We performed whole-cell recordings of excitatory neurons in the forelimb region of M1 (Fig. 1A) to assess the effect of impaired dopamine signaling locally and/or across the motor circuit on the input/output function of pyramidal neurons in superficial and deep layers of M1. Recorded neurons included in this study showed pyramidal morphology and were negative for GAD67 immunoreactivity (Fig. 1B). In order to determine laminar borders, sections containing M1 were stained for the cytoarchitectural marker SMI-32, a nuclear counterstain Hoechst33342, and a fluorescent Nissl (Fig. 1C). The top border of Layer 2/3 (L23) was placed where cell density sharply drops off as you move towards the pial surface. The bottom of L2/3 was defined at the depth where the cortex transitions from small, densely packed neurons to very large, more sparse pyramidal neurons, as visualized in the Nissl. SMI-32 labels a subset of pyramidal neurons in L2/3 and Layer 5 (L5) (Campbell and Morrison, 1989; Voelker et al., 2004), and in dysgranular and agranular cortex it is expressed most strongly in L5 (Barbas and Garcia-Cabezas, 2016). Staining for SMI-32 was used to confirm the previously defined laminar borders, and to demarcate the end of L5 and the beginning of Layer 6. Cells were localized to L2/3 or L5 by determining their depth from the cortical surface with *post hoc* immunostaining of recorded neurons.

**Figure 1.**
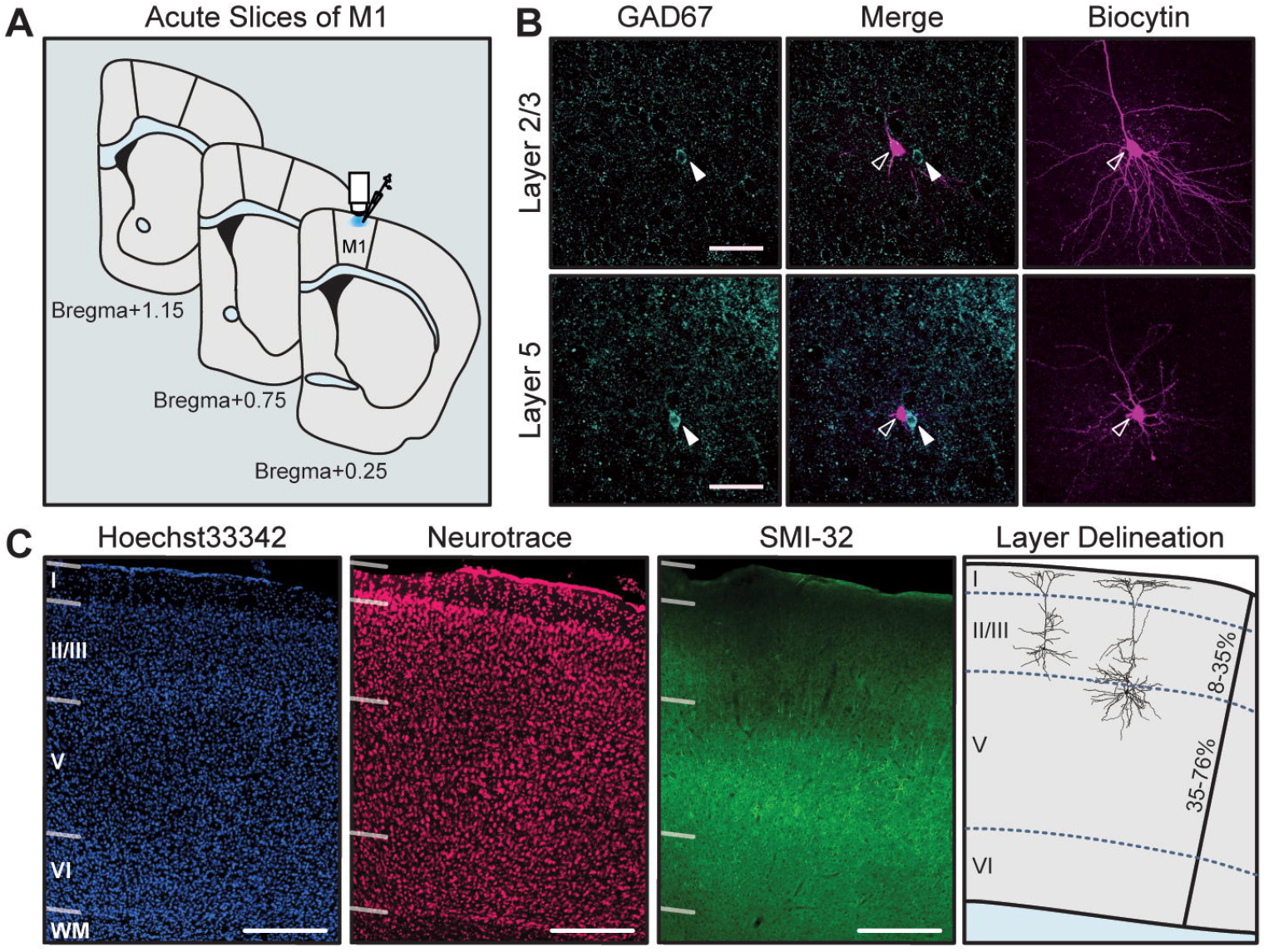
Whole-cell recordings of excitatory neurons in forelimb M1, localized to L2/3 and L5. **(A)** Schematic showing the anterior-posterior span of recorded slices, restricted to forelimb area of M1. **(B)** Recorded neurons were visualized with streptavidin labeling of biocytin and confirmed as excitatory by negative immunoreactivity for GAD67. GAD67 and Merge images shown at one z-plane depth; biocytin images shown as a collapsed stack spanning the entire neuron. Open arrows: GAD67^−^ biocytin-filled neurons; closed arrows: GAD67^+^ interneurons (not recorded) at same depth. Scale bar = 50μm. **(C)** Histological staining of cytoarchitecture used to define cortical layers. Hoechst33342 is a nuclear counterstain of all cells in the region, Neurotrace was used as a neuron-specific stain for somata, SMI-32 labels a subset of pyramidal neurons in layer 3 and layer 5. Scale bar = 200 μm. Rightmost panel: two example neurons localized to L2/3 and L5: neurons localized within 8-35% of the total cortical depth were defined as L2/3; neurons within 35-76% of cortical depth were defined as L5.

### Acute Blockade of Dopamine Receptors Increases M1 Neural Excitability

D1R and D2R in the rodent brain are expressed in neurons across the cortical mantle, although their expressions show laminar preference (Lemberger et al., 2007; Santana et al., 2009). While dopaminergic modulation through these receptors influences activity in M1 (Molina-Luna et al., 2009; Hosp et al., 2011), the mechanisms underlying dopamine’s regulation of M1 neurons are not clear. To distinguish how D1R and D2R modulate the input/output function of M1 neurons, we first studied the effects of acute dopamine receptor blockade using pharmacological antagonists. We compared current clamp responses to subthreshold and suprathreshold current steps before and after bath application of dopamine receptor antagonists. To determine whether the modulation of pyramidal neuron input/output function by dopamine receptors is due to intrinsic conductance or synaptic activity, we compared the effect of dopamine receptor antagonists in ACSF, in which spontaneous synaptic activity is present, and in the presence of ionotropic GABA and glutamate receptor blockers.

In a first set of experiments, we examined the effects of the D1R antagonist SCH23390 (10 μM; D1ant) on the excitability (see Methods for detailed description of analysis) of pyramidal neurons in L2/3 and L5 of M1 (Fig. 2). In ACSF, bath application of SCH23290 increased the dynamic input resistance of both L2/3 and L5 neurons (Fig. 2B; in MΩ: L2/3 ACSF = 141.32±9.97, L2/3 D1ant = 170.49±12.24, p=0.007; L5 ACSF = 106.33±14.80, L5 D1ant = 123.28±15.66, p=0.007). This effect was potentiated in the presence of ionotropic GABA and glutamate receptor blockers (BLK), suggesting that increased dynamic input resistance induced by D1R blockade occurs independent of fast synaptic transmission (Fig. 2B, in MΩ: L2/3 BLK = 139.06±12.04, L2/3 D1ant = 201.73±20.41, p=0.001; L5 BLK = 91.79±11.29, L5 D1ant = 136.25±20.90, p=0.002).

**Figure 2.**
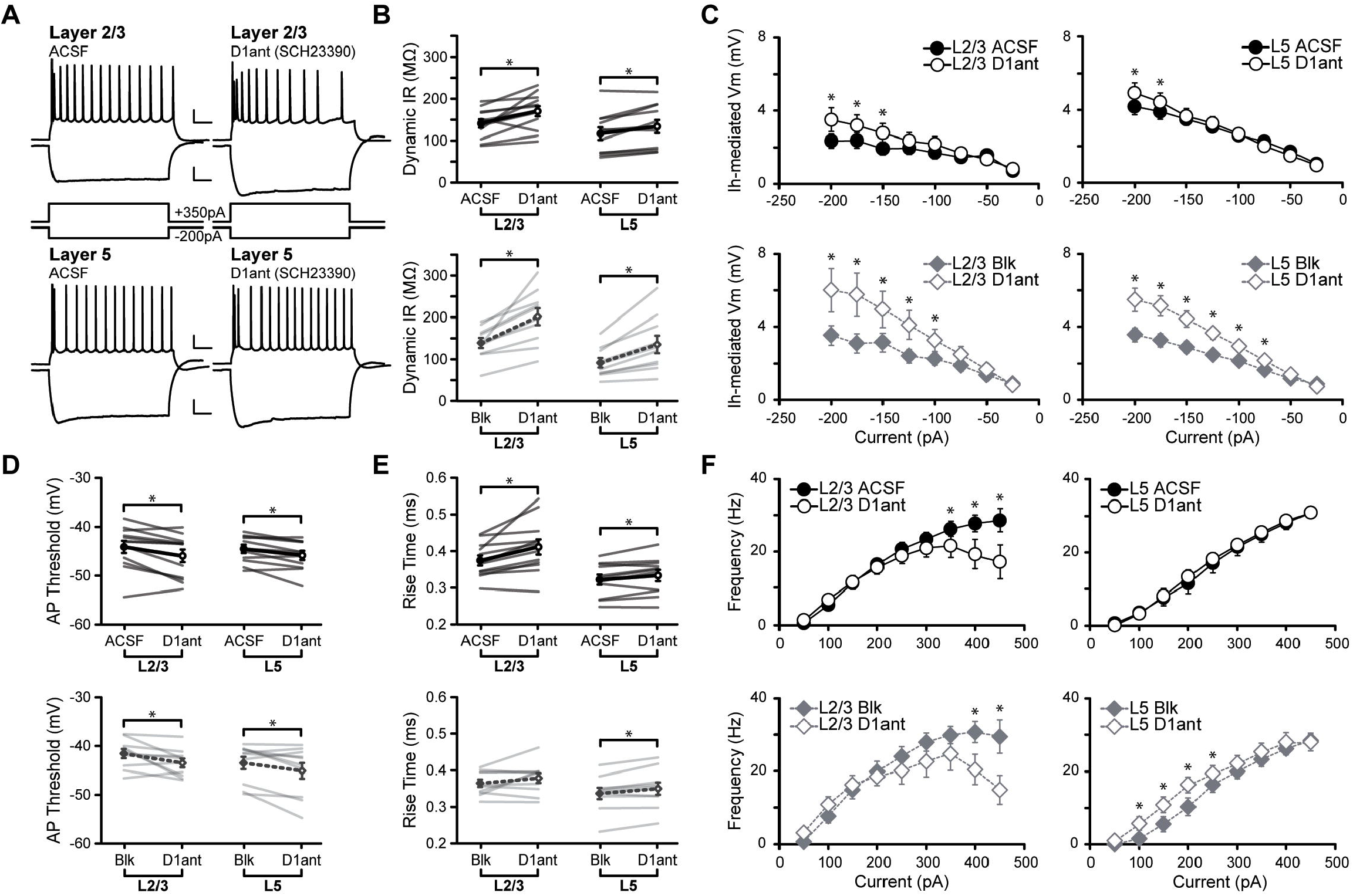
Acute D1R blockade shifts excitability of M1 neurons. **(A)** Superimposed responses to hyperpolarizing and depolarizing current steps in individual L2/3 and L5 neurons before and after bath application of D1R antagonist SCH23390 (D1ant: 10μM), upper scale bars: 20mV, 100ms, lower scale bars: 10mV, 100ms. **(B-F)** Summary excitability plots for excitatory neurons in L2/3 and L5 before and after D1ant application, in baseline (ACSF) or synaptic blocker (Blk: 20μM picrotoxin, 20μM DNQX, 50μM AP5) conditions. **(B)** Dynamic input resistance across hyperpolarizing current steps. **(C)** The amount of I_h_-mediated voltage sag elicited by hyperpolarizing current. **(D)** Action potential threshold at rheobase. **(E)** Action potential rise time at rheobase. **(F)** Action potential frequency during suprathreshold current injections. (ACSF L2/3 neurons N=6 n=12, ACSF L5 neurons N=6 n=11, Blk L2/3 neurons N=5 n=10, Blk L5 neurons N=6, n=10; data shown as mean±SEM; *denotes p<0.05).

In response to hyperpolarizing current steps, neurons showed a voltage sag during the initial portion of the response, which is typically associated with the presence of hyperpolarization-activated cation channels (HCN) mediating I_h_ (Rosenkranz and Johnston, 2006; Hogan and Poroli, 2008). Bath application of SCH23390 increased the amplitude of the voltage sag in both L2/3 and L5 neurons, and this change persisted in the presence of synaptic blockers, confirming the independence of this effect from fast synaptic transmission (Fig. 2C).

Next, we assess the effect of D1R blockade on the suprathreshold portion of the input/output function. We first compared action potential properties at rheobase: The action potential threshold was more hyperpolarized in both L2/3 and L5 neurons in the presence of SCH23390 (Fig. 2D; in mV: L2/3 ACSF = −44.10±1.26, L2/3 D1ant = −45.96±1.28, p=0.0073; L5 ACSF = −44.57±0.84, L5 D1ant = −45.87±0.93, p=0.0087). When this experiment was repeated in the presence of fast synaptic transmission blockers, the effect remained in both layers (Fig. 2D, in mV: L2/3 BLK = −41.57±0.93, L2/3 D1ant = −43.47±0.88, p=0.048; L5 BLK = −43.49±1.25, L5 D1ant = −45.14±1.63, p=0.047). Blocking D1Rs also increased action potential rise time in both L2/3 and L5 neurons (Fig. 2E, in ms: L2/3 ACSF = 0.37±0.01, L2/3 D1ant = 0.41±0.02, p=0.0069; L5 ACSF = 0.32±0.01, L5 D1ant = 0.33±0.01, p=0.033). This effect of D1R blockade was eliminated in L2/3 but persisted L5 neurons in the presence of synaptic transmission blockers (Fig. 2E, in ms: L2/3 BLK = 0.37±0.01, L2/3 D1ant = 0.38±0.01, p=0.18; L5 BLK = 0.34±0.02, L5 D1ant = 0.35±0.02, p=0.01). Thus, dopamine affects action potential properties of M1 pyramidal neurons by acting through distinct mechanisms in superficial and deep layers.

Bath application of the D1R antagonist SCH23390 unveiled laminar differences in the effects of acute D1R blockade on the input/output curve. Comparison of relationships between action potential frequency versus injected current (fI curve) in ACSF and acute D1R blockade showed that the ability of L2/3 neurons to increase their firing rates in response to increasing current steps was impaired, significantly reducing the maximum firing rate. In contrast, SCH23390 did not affect the fI curve of L5 neurons (Fig. 2F). The changes in the fI curve of L2/3 neurons persisted in synaptic blockers, indicating that it depends on modulation of voltage gated conductance. Interestingly, in L5, pharmacological blockade of ionotropic synaptic receptors unveiled a previously masked effect of D1R blockade on the fI curve: an increase in firing rate selectively in the linear portion of the fI curve, the range in which neurons’ firing rates show high sensitivity to small changes in current injections. The increase in input resistance and hyperpolarization in action potential threshold point to a net increase in M1 pyramidal neuron excitability in the absence of dopamine activation of D1R. In L2/3, selectively blocking dopamine signaling through D1R also results in a decreased maximum firing rate, suggesting that large incoming input would result in reduced output.

Dopaminergic modulation of neuronal activity can also rely on D2Rs, which are also expressed in M1 neurons (Santana et al., 2009). Previous *in vivo* studies reported that activation of D2Rs increases the firing rate of M1 pyramidal neurons in anesthetized animals and can alter motor maps, but the mechanisms underlying these effects are unclear (Hosp et al., 2009; Vitrac et al., 2014). To assess whether D2Rs modulate the input/output curve of M1 pyramidal neurons we repeated the experiments above using the D2R antagonist sulpiride (10μM; Fig. 3; D2ant). Bath application of sulpiride increased the input resistance of L2/3 and showed a trend toward increased input resistance in L5 neurons (Fig. 3B, in MΩ: L2/3 ACSF = 108.56±11.01, L2/3 D2ant = 127.78±14.12, p=0.019; L5 ACSF = 97.1763±11.94, L5 D2ant = 107.09±12.69, p=0.06). The increase in input in L2/3 neurons was eliminated by GABAA, AMPA and NMDA receptor antagonists, suggesting that it relies on modulation of synaptic transmission. In contrast, application of sulpiride in L5 in the presence of synaptic receptor blockers amplified the increase in input resistance (Fig. 3B, in MΩ: L2/3 BLK = 137.33±25.23, L2/3 D2ant = 148.17±28.67, p=0.13; L5 BLK = 78.69±11.69, L5 D2ant = 95.63±17.39, p=0.027). Sulpiride did not affect the voltage sag in either L2/3 or L5 pyramidal neurons (Fig. 3C), suggesting that D2R do not modulate this current in M1.

**Figure 3.**
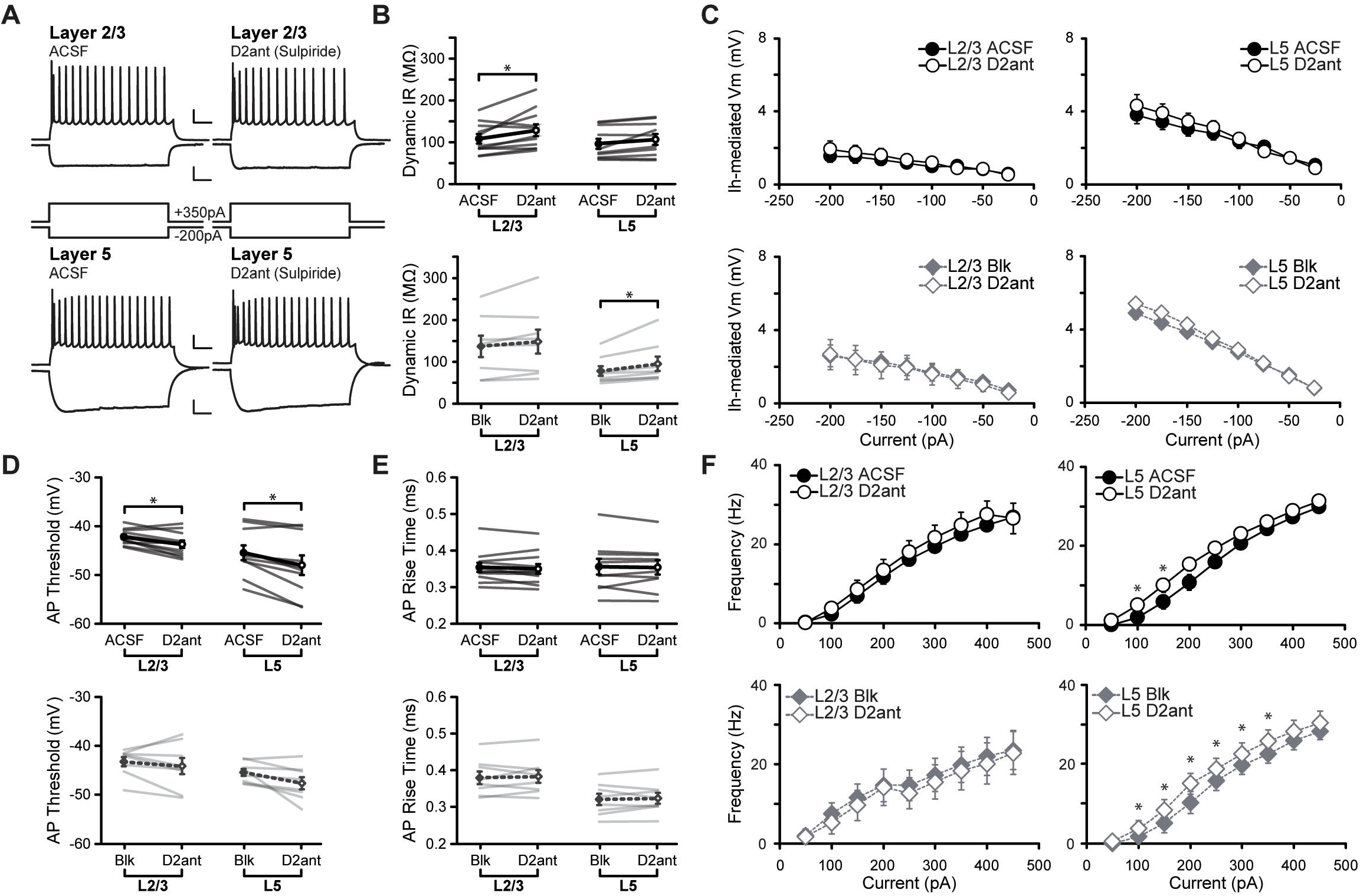
Acute DR2 blockade shifts excitability of M1 neurons. **(A)** Superimposed responses to hyperpolarizing and depolarizing current steps in individual L2/3 and L5 neurons before and after bath application of D2R antagonist Sulpiride (D2ant: 10μM), upper scale bars: 20mV, 100ms, lower scale bars: 10mV, 100ms. **(B-F)** Summary excitability plots for excitatory neurons in L2/3 and L5 before and after D2ant application, in baseline (ACSF) or synaptic blocker (Blk: 20μM picrotoxin, 20μM DNQX, 50μM AP5) conditions. **(B)** Dynamic input resistance across hyperpolarizing current steps. **(C)** The amount of I_h_-mediated voltage sag elicited by hyperpolarizing current. **(D)** Action potential threshold at rheobase. **(E)** Action potential rise time at rheobase. **(F)** Action potential frequency during suprathreshold current injections. (ACSF L2/3 neurons N=5 n=11, ACSF L5 neurons N=5 n=10, Blk L2/3 neurons N=4 n=8, Blk L5 neurons N=4, n=8; data shown as mean±SEM; *denotes p<0.05)

**Figure 4.**
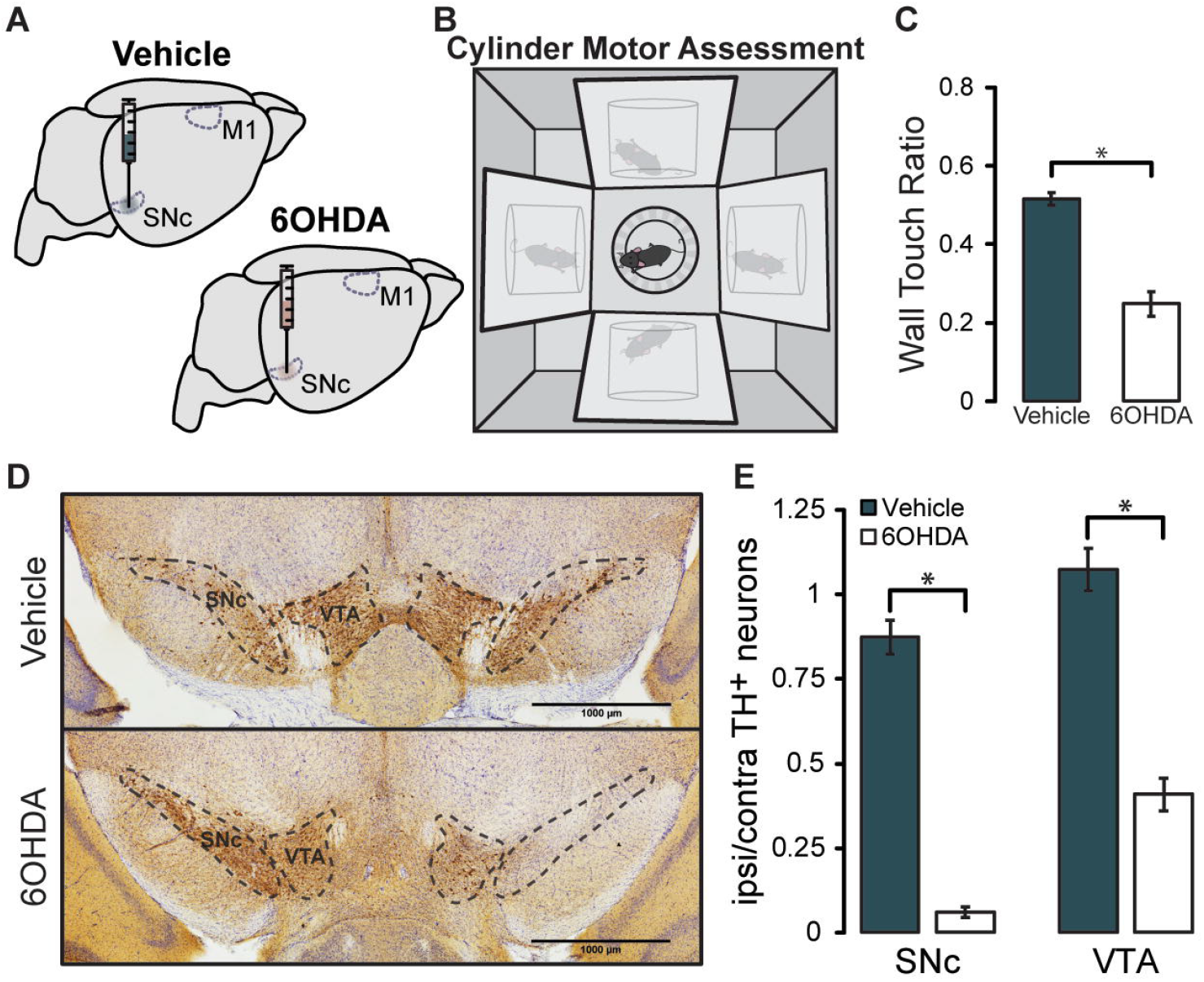
Validation of 6OHDA Model of Parkinson’s Disease. **(A)** Unilateral injection of 6OHDA, or vehicle, centered on the SNc. **(B)** Schematic of the cylinder motor assessment. **(C)** Quantification of weight-bearing wall touches measured as a ratio of forelimb use contralateral versus ipsilateral to the injected hemisphere. **(D)** Immunolabeled TH^+^ dopaminergic neurons visualized with DAB in the SNc and VTA. **(E)** Summary of stereological counts of TH^+^ neurons in the SNc and VTA of lesioned or vehicle-injected animals. (Vehicle Animals N=12, 6OHDA Animals N=8, data shown as mean±SEM; *denotes p<0.0001).

Bath application of sulpiride hyperpolarized the action potential threshold (Fig. 3D, in mV: L2/3 ACSF = −42.22±0.49, L2/3 D2ant = −43.72±0.75, p=0.0063; L5 ACSF = −45.46±1.51, L5 D2ant = −48.05±2.00, p=0.003). In both layers this effect was eliminated by in the presence of ionotropic GABA and glutamate receptor blockers (Fig. 3D, in mV: L2/3 BLK = −43.26±0.98, L2/3 D2ant = −44.15±1.68, p=0.41; L5 BLK = −45.48±0.75, L5 D2ant = −47.72±1.25, p=0.054). Sulpiride had no effect on action potential rise time (Fig. 3E, in ms: L2/3 ACSF = 0.35±0.01, L2/3 D2ant = 0.35±0.01, p=0.32; L5 ACSF = 0.36±0.02, L5 D2ant = 0.35±0.02, p=0.69; L2/3 BLK = 0.38±0.02, L2/3 D2ant = 0.38±0.02, p=0.65; L5 BLK = 0.32±0.02, L5 D2ant = 0.32±0.01, p=0.63). Finally, sulpiride had no effect on the fI curve of L2/3 pyramidal neurons. However, in L5 it increased the action potential frequency in the linear portion of the fI curve, increasing L5 neurons output to small changes in input current. The effect of sulpiride on the fI curve of L5 neurons persisted in the presence of synaptic blockers, suggesting that this effect is independent of fast synaptic transmission (Fig. 3F). Taken together, these data suggest that while D2R play a role in M1 excitability, the consequences of an acute loss of D2R activity is more subtle than that of D1R antagonism. In both layers, acute blockade of D2R signaling results in a net increase in excitability. While in L2/3 this effect depends on synaptic transmission, in L5 is primarily due to modulation of intrinsic conductance.

Taken together, these results point to laminar specific effects of D1R and D2R activity in M1, and support the hypothesis that dopamine directly modulates the excitability of M1 pyramidal cells through a complex set of mechanisms that involve intrinsic conductance and synaptic transmission. D1R and D2R have partial, but not completely overlapping effects on the input/output function of M1 pyramidal neurons.

### Chronic Midbrain Dopamine Depletion Alters M1 Neural Excitability

Diminished dopamine signaling in the motor system and progressive motor impairment are hallmarks of PD. Patients, as well as animal models of the disease, exhibit motor cortex dysfunction. We hypothesized that chronic loss of dopaminergic input to the motor circuit alters the excitability of M1 pyramidal neurons, possibly providing a mechanism for impaired motor cortex activity. We asked whether chronic loss of dopaminergic activity across the entire motor circuit, or locally within M1, is sufficient to shift the excitability of M1 neurons and recapitulate the results of the acute dopamine receptor blockade experiments. We unilaterally injected 6OHDA, or an equivalent volume of vehicle as a control, in the midbrain centering the injection site on the SNc (Fig. 4). Dopamine depletion of midbrain dopaminergic neurons with 6OHDA is widely used as a model of PD and is known to induce motor impairment. Two weeks after injection, and just prior to recording, each animal’s movement was assessed with a cylinder motor task (Fig. 4B). 6OHDA-injected mice showed reduced use of the forelimb contralateral to the injection, while vehicle injected animals showed no sign of forelimb use impairment (Fig. 4C, Vehicle = 0.52±0.02, 6OHDA = 0.25±0.03, p=3.58×10^−8^). The severity of the 6OHDA lesion was anatomically assessed with *post hoc* immunostaining for tyrosine hydroxylase (TH), followed by unbiased stereological counts of TH^+^ neurons in the VTA and SNc. Lesioned animals showed significant cell loss in both the SNc and VTA when compared to their vehicle-injected counterparts (Fig. 4D, E, ipsi/contra TH^+^ neurons, Vehicle SNc = 0.87±0.03, 6OHDA SNc = 0.06±0.03, p=1.6×10^−12^; Vehicle VTA = 1.07±0.07, 6OHDA VTA = 0.41±0.03, p=5.3×10^−7^), indicating that the unilateral injection had effectively depleted dopamine neurons and induced the expected motor impairment.

We then assessed the effect of the 6OHDA manipulation on the input/output function of L2/3 and L5 pyramidal neurons in M1 (Fig. 5). In contrast to the acute blockade of D1R and D2R, chronic midbrain dopamine depletion did not alter input resistance or action potential threshold of either L2/3 or L5 neurons in M1 (Fig. 5B, D). Interestingly, the voltage sag was unaffected in L2/3 neurons, but significantly reduced in L5 (Fig. 5C), which opposes the effect we observed following acute D1R antagonism. Further, this effect was eliminated when the experiment was performed in synaptic blockers, pointing to the involvement of synaptic mechanisms in this result.

**Figure 5.**
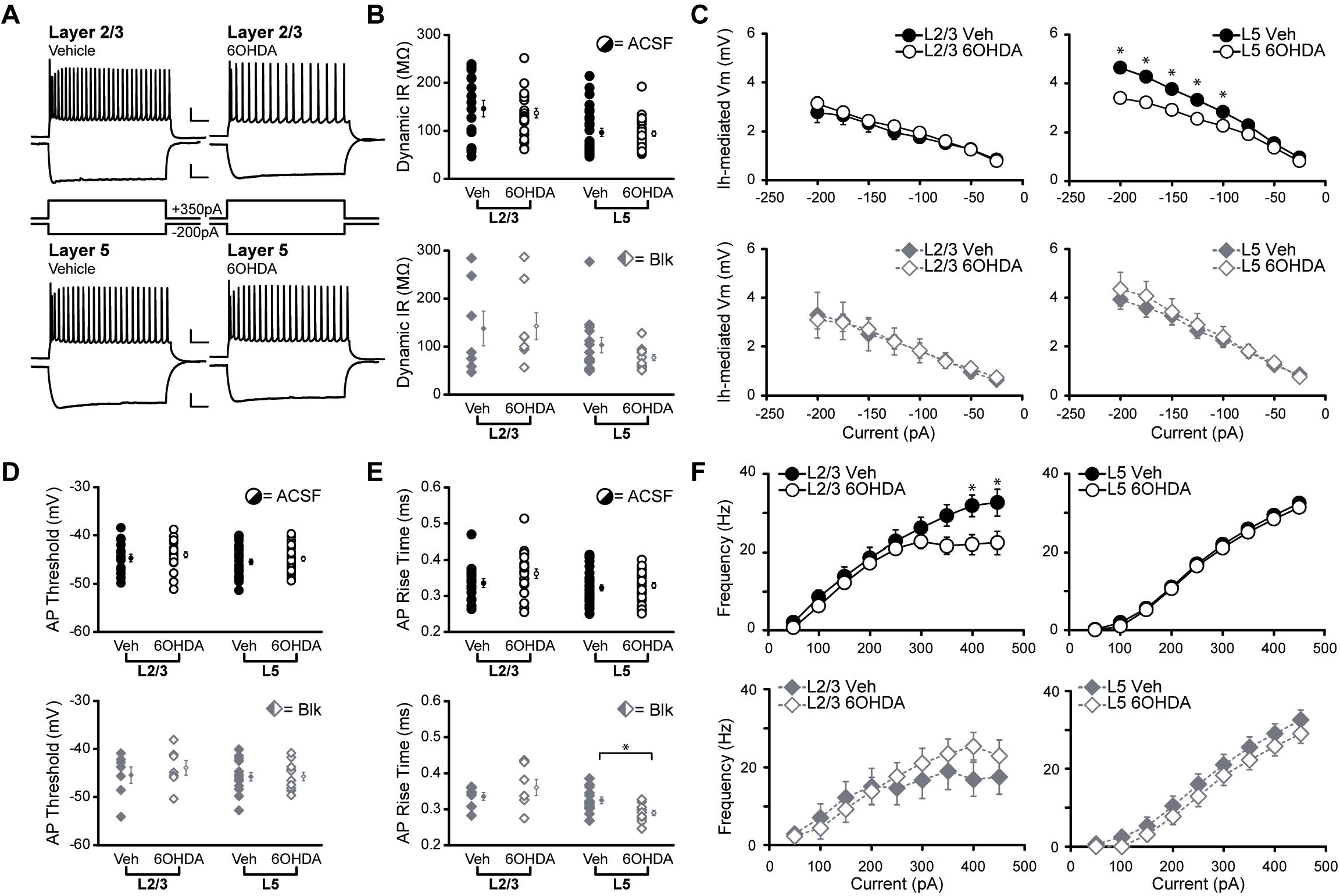
Nigral 6OHDA lesion shifts M1 neuron excitability, partially recapitulating effects of D1R antagonist. **(A)** Superimposed responses to hyperpolarizing and depolarizing current steps in individual L2/3 and L5 neurons of vehicle and 6OHDA-injected animals. All traces shown are in ACSF conditions. Upper scale bars: 20mV, 100ms, lower scale bars: 10mV, 100ms. **(B-F)** Summary excitability plots for excitatory neurons in L2/3 and L5 of vehicle and 6OHDA animals, in baseline (ACSF) or synaptic blocker (Blk: 20μM picrotoxin, 20μM DNQX, 50μM AP5) conditions. **(B)** Dynamic input resistance across hyperpolarizing current steps. **(C)** The amount of I_h_-mediated voltage sag elicited by hyperpolarizing current. **(D)** Action potential threshold at rheobase. **(E)** Action potential rise time at rheobase. **(F)** Action potential frequency during suprathreshold current injections. (ACSF L2/3 vehicle neurons N=11 n=16, ACSF L2/3 6OHDA neurons N=13 n=21, Blk L2/3 vehicle neurons N=3 n=7, Blk L2/3 6OHDA neurons N=4 n=8, ACSF L5 vehicle neurons N=11, n=29, ACSF L5 6OHDA neurons N=11 n=29, Blk L5 vehicle neurons N=5 n=14, Blk L5 6OHDA neurons N=6 n=12; data shown as mean±SEM; *denotes p<0.05).

Action potential rise time was not significantly altered in ACSF, however blockade of fast synaptic transmission unveiled a reduction in this parameter specifically in L5 pyramidal neurons (Fig. 5E, in ms: L2/3 Vehicle BLK = 0.34±0.01, L2/3 6OHDA BLK = 0.36±0.02, p=0.33; L5 Vehicle BLK = 0.32±0.01, L5 6OHDA BLK = 0.29±0.01, p=0.0088). This result suggests that the effect of 6OHDA relies on changes in intrinsic properties and was masked by synaptic transmission.

Analysis of fI curves for both L2/3 and L5 pyramidal neurons unveiled an effect similar to the effect of acute D1R blockade: L2/3 neurons of 6OHDA mice lost the ability to increase their firing rate at the highest current injections (Fig. 5F), and this effect was abolished by application of synaptic blockers. The fI curve of L5 neurons in 6OHDA mice was not significantly different from that of vehicle injected mice both in either ACSF or synaptic blockers (Fig. 5F). These results suggest that midbrain depletion of dopamine induces laminar specific changes in M1 pyramidal neurons excitability, likely through a combination of altered dopaminergic signaling in M1 as well as a shift in overall synaptic transmission in the area.

### Chronic depletion of dopaminergic input to M1 exclusively impacts L2/3 excitability

The partial overlap in results between the acute dopamine receptor antagonism and chronic midbrain dopamine depletion led us to ask if reduced, chronic dopaminergic innervation within M1 could be a contributing factor. M1 receives a direct dopaminergic projection from midbrain regions, with the majority of this input originating in the VTA as well as a small input from the SNc (Hosp et al., 2011; Leemburg et al., 2018). Our 6OHDA lesion affected primarily the SNc but also reduced the number of dopaminergic neurons in the VTA. We therefore tested if the effects of midbrain dopamine depletion could be recapitulated by a 6OHDA lesion within M1 (Fig. 6).

**Figure 6.**
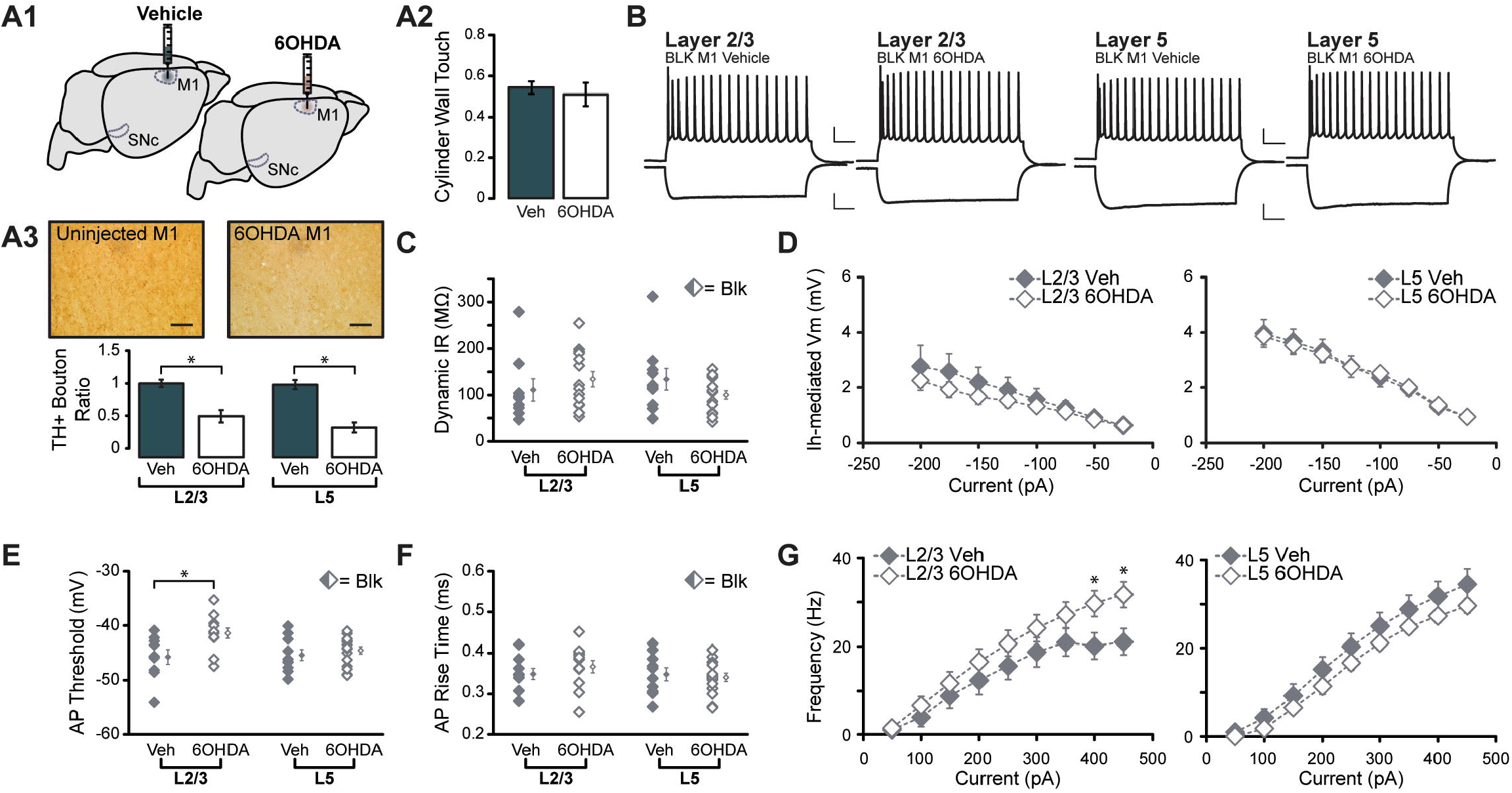
Chronic M1 dopamine depletion impacts L2/3 intrinsic excitability. **(A1)** Unilateral injection of 6OHDA, or vehicle, into forelimb region of M1. **(A2)** M1 dopamine depletion was not sufficient to influence lesioned animal performance in cylinder motor assessment. **(A3)** Top panel: TH+ axons and boutons labeled in L2/3 of M1 ipsilateral and contralateral to the injection site (magnification: 40x, scale bar: 50μm). Bottom panel: The ratio of TH+ boutons in L2/3 and L5 between injected and uninjected hemispheres was reduced in 6OHDA animals (L2/3 p<0.01, L5 p<0.001). **(B-F)** Summary excitability plots for excitatory neurons in L2/3 and L5 of vehicle or 6OHDA-injected animals, performed in synaptic blockers (Blk: 20μM picrotoxin, 20μM DNQX, 50μM AP5). **(B)** Superimposed responses to hyperpolarizing and depolarizing current steps in individual L2/3 and L5 neurons in vehicle or 6OHDA-injected animals, upper scale bars: 20mV, 100ms, lower scale bars: 10mV, 100ms. **(C)** Dynamic input resistance across hyperpolarizing current steps. **(D)** The amount of I_h_-mediated voltage sag elicited by hyperpolarizing current. **(E)** Action potential threshold at rheobase. **(F)** Action potential rise time at rheobase. **(G)** Action potential frequency during suprathreshold current injections. (Vehicle L2/3 neurons N=5 n=9, Vehicle L5 neurons N=5 n=10, 6OHDA L2/3 neurons N=4 n=14, 6OHDA L5 neurons N=4, n=16; data shown as mean±SEM; *denotes p<0.05).

For these experiments, a unilateral 6OHDA lesion, or a vehicle injection, was centered on the forelimb region of M1. Two weeks after injection, mice were tested for motor impairment in the cylinder task described above. 6OHDA lesions in M1 did not induce motor impairment in the cylinder motor task compared to vehicle injected controls (Fig. 6A2, Vehicle = 0.54±0.03, 6OHDA = 0.51±0.06, p=0.6). Stereological counts of TH^+^ boutons in L2/3 and L5 of M1 indicated a significant reduction of TH^+^ boutons in both layers of 6OHDA-injected M1 (Fig. 6A3, Vehicle L2/3 = 1.13±0.06, 6OHDA L2/3 0.63±0.10, p=0.019; Vehicle L5 = 1.12±0.07, 6OHDA L5 = 0.46±0.08, p=0.00041), indicating that the 6OHDA injection was effective.

As the effects of acute D1R and D2R blockade were independent of synaptic transmission, patch clamp recordings in M1 lesioned mice were carried out in the presence of synaptic blockers. L5 pyramidal neurons showed no differences in the subthreshold or suprathreshold range of activity (Fig. 6C-G), suggesting that chronic loss of dopaminergic input to M1 did not affect the excitability of these neurons. In contrast, L2/3 neurons showed two major changes. First, the action potential threshold was significantly depolarized compared to vehicle injected mice (Fig. 6E, in mV: L2/3 Vehicle = −45.76±1.36, L2/3 6OHDA = −41.39±0.93, p=0.013), indicating larger currents are required for these neurons to generate an action potential. However, the firing frequency in response to suprathreshold current injections was increased compared to vehicle injected mice (Fig. 6G), suggesting that the output of these neurons in response to larger inputs is increased. These shifts in L2/3 neuron excitability are unique to the M1 lesion experiment and are not recapitulated by either dopamine receptor blockade or midbrain dopaminergic cell loss. This suggests that local dopaminergic deafferentation within M1 causes unique changes in excitability specific to L2/3 pyramidal neurons. Taken together, our results indicate that dopamine impairment can have complex effects on the input/output function of M1 neurons depending on the duration and location of the dopamine impairment.

## Discussion

Dopaminergic signaling is crucial for skilled voluntary movement, and reduced dopamine in the motor circuit leads to motor impairment. PD is characterized by progressive dopaminergic cell death in the SNc, which primarily projects to the basal ganglia, and a less severe but significant loss of dopaminergic neurons in the VTA (Giguere et al., 2018). While the effects of reduced dopamine signaling has been well-documented in the basal ganglia (Blandini et al., 2000; Day et al., 2006; Azdad et al., 2009; Bagetta et al., 2010; Fieblinger et al., 2014), how midbrain dopaminergic cell death affects M1 is less clear. Previous studies have highlighted the importance of dopamine in M1 for motor learning and plasticity (Molina-Luna et al., 2009; Rioult-Pedotti et al., 2015), and impairment of dopaminergic input to M1 *in vivo* results in impaired skill learning, delayed movement execution, and structural changes at M1 synapses (Hosp et al., 2011; Guo et al., 2015; Chen et al., 2019), however the mechanisms underlying these changes are understudied. We demonstrate that impaired dopamine signaling impacts the excitability of M1 pyramidal neurons and could be contributing to diminished motor function observed in previous studies.

Acute D1R and D2R antagonism impacts the input/output function of M1 neurons, suggesting that dopamine regulates excitability in a healthy M1. Our results show that D1R blockade increases excitability of L2/3 and L5 neurons in the subthreshold range of activity up to rheobase. Within the suprathreshold range of activity, on the other hand, layer-specific effects emerge. The fI curve of L2/3 neurons was shifted downward following D1R antagonism, particularly impacting the response to large input. In contrast, L5 neurons showed an upwards shift in the linear portion of the fI curve when fast synaptic transmission was blocked. This implies that D1R antagonism caused increased excitability of L5 neurons in their suprathreshold range as well, however this effect was masked when spontaneous synaptic transmission was present. In fact, almost all effects of D1R antagonism persisted in the presence of fast synaptic transmission blockers, consistent with direct expression of these receptors on the cells we recorded and with the engagement of intrinsic mechanisms for the modulation of membrane properties. The effects of D2R blockade were more subtle, although consistent with an overall increase in excitability in L2/3 and L5. Increased input resistance and hyperpolarization of action potential threshold were eliminated by synaptic blockers, indicating that they depend on modulation of incoming synaptic drive and not intrinsic properties. Selectively in L5, D2R blockade caused an upward shift in the linear portion of the fI curve. This effect was amplified by synaptic transmission blockade, pointing to a direct effect of D2R on L5 pyramidal neurons excitability. These data indicate that, in M1, D2Rs may have a more restricted expression than D1R, or that their primary role is modulating synaptic transmission.

The increase in excitability following D1R blockade is surprising in view of studies in the basal ganglia showing that dopamine or D1R agonists typically increase excitability (Planert et al., 2013). However, reports show that dopamine, via D1R, reduces excitability of L5 pyramidal neurons in the entorhinal cortex (Rosenkranz and Johnston, 2006). Additionally, dopamine application in M1 *in vivo* reduces spontaneous firing of corticospinal neurons (Huda et al., 2001; Awenowicz and Porter, 2002). Our results are consistent with these findings, particularly in L5, where acute dopamine receptor blockade increased excitability in a layer known to project to the pyramidal tract (Oswald et al., 2013).

After establishing that the excitability of M1 pyramidal neurons is directly modulated by dopamine, we extended our study to assess how chronic loss of dopamine modulates the excitability of M1 pyramidal neurons. Loss of dopaminergic neurons projecting to the motor circuit, particularly to the basal ganglia, leads to movement dysfunction and is a hallmark of PD. Chronic depletion of dopamine impacts neural activity across the motor circuit (Blandini et al., 2000; Day et al., 2006; Azdad et al., 2009; Planert et al., 2013; Benazzouz et al., 2014), and we posited that chronic loss of dopamine to all areas involved in motor control would lead to reverberating changes in excitability within M1, despite this region receiving only a fraction of the dopaminergic input. In agreement with previous work, our 6OHDA-injected mice showed unilaterally impaired forelimb use. Furthermore, 6OHDA mice showed laminar-specific shifts in excitability within M1, indicating that neurons in L2/3 and L5 have distinct sensitivity to chronic loss of dopamine signaling. Contrasting the effects of acute D1R blockade, L2/3 neurons in 6OHDA mice showed no change in the subthreshold range of activity. However, the fI curve of L2/3 neurons showed impaired response to large inputs, mirroring the effect of acute D1R antagonism. This change was eliminated by synaptic transmission blockers, suggesting that unlike the effect of acute D1R blockade, this comparable shift in the fI curve of L2/3 neurons is dependent on synaptic transmission. The excitability of L5 neurons was also affected by chronic dopamine depletion, although the effects were unique to this cortical layer and dopamine manipulation. Chronic midbrain dopamine depletion induced a decrease in I_h_ of L5 cells with no changes in the suprathreshold range of their input/output function. The decrease in I_h_ was fully eliminated by application of fast synaptic transmission blockers, suggesting it is mediated by synaptic activity. Decreased I_h_ alters the capacity of the neuron to maintain resting membrane potential and impacts the response to input, particularly those that would hyperpolarize the cell.

Overall, the changes in excitability due to midbrain dopamine loss rely on synaptic drive, suggesting that chronic midbrain dopamine manipulation may alter synaptic transmission into or within M1. This could occur as a consequence of shifting activity within the basal ganglia following dopamine loss, as an effect of reduced dopamine signaling directly in M1, or a combination of these factors. While a thorough analysis of synaptic transmission is beyond the scope of this study, there are reports of altered synaptic activity in models of PD elsewhere in the motor circuit (Day et al., 2006; Fan et al., 2012; Galvan et al., 2015; Parker et al., 2016), leading to a hypothesis that excitatory drive of M1 may be reduced.

We predicted that changes to M1 pyramidal neuron input/output function following midbrain dopamine depletion would likely arise from altered activity across the motor circuit, rather than simply from loss of dopaminergic input exclusively to M1. In support of this interpretation, restricting the 6OHDA injection to M1 altered the excitability of L2/3, but not L5, pyramidal neurons with apparently opposing effects to those caused by midbrain dopamine depletion. The action potential threshold of L2/3 pyramidal neurons was depolarized, indicating a decrease in excitability in the subthreshold range of activity. When these cells were driven to fire, they responded with a higher action potential frequency than those recorded in vehicle-injected mice, suggesting an increase in excitability in the suprathreshold range of activity. Such effects may depend on modulation of distinct conductance regulating different aspects of action potential dynamics. These results differ from those obtained following all other manipulations, and we speculate that they result from a combinatorial, chronic reduction in M1 D1R and D2R activity, or may result from compensatory mechanisms following local loss of dopaminergic innervation. This manipulation did not induce motor impairment, pointing to a limited impact of direct dopaminergic projections in motor execution. This result is expected, as direct dopamine modulation in M1 is primarily associated with synaptic plasticity and motor learning, aspects that we did not examine (Molina-Luna et al., 2009; Hosp et al., 2011).

### Functional Implications

Dopamine receptor blockade increased the excitability of L2/3 and L5 neurons. In M1, L2/3 pyramidal neurons are mainly corticocortical or corticostriatal-projecting, while those in L5 are corticospinal, corticothalamic, and corticostriatal-projecting (Oswald et al., 2013). Signals flow superficial to deep, with high intracortical connectivity between L2/3 neurons and corticospinal neurons in L5 (Weiler et al., 2008). Our results show acute impairment of D1R and D2R signaling increases the excitability of L5 neurons and that of one of their primary presynaptic partners. Together, these changes may lead to hyperactivity in M1, an effect associated with motor impairment and movement disorders like PD (Thobois et al., 2000; Haslinger et al., 2001). Our results also show that dopamine receptor antagonism increased the I_h_-mediated sag in L2/3 and L5, while midbrain 6OHDA injections decreased the sag selectively in L5. In M1, I_h_ is thought to be a regulator of signal flow from superficial layers, involved in motor planning, to deeper output layers (Sheets et al., 2011). We show that dopamine can directly and indirectly influence the activity of I_h_, providing evidence for one source of neuromodulatory control of signal flow in M1. Interestingly, one study of an animal model of PD reported downregulation of HCN2 channels, which in part mediate I_h_, in the globus pallidus (Chan et al., 2011). This led to abolished autonomous pacemaking activity and induced abnormal synchronous activity. While re-introduction of HCN2 channels in the globus pallidus restored normal signaling, it was not sufficient to recover motor impairments. In view of our findings, one may speculate that abnormal HCN activity in PD may extend beyond the globus pallidus, and that coordinated restoration of the activity of these channels may be required for symptom improvement.

### Conclusions

Dopamine signaling in the motor system is crucial for the execution of voluntary movements. While most work focuses on the effects of dopamine signaling in the basal ganglia, recent studies point to M1 as an additional site of dysfunction in PD patients and mouse models of the disease. Our results indicate that diminished dopamine signaling, whether acute or chronic, has profound effects on the excitability of M1 neurons. We unveil a complex combination of laminar specific mechanisms for dopamine-dependent modulation of pyramidal neuron excitability, which are likely to significantly alter the output of M1 and influence movement execution.

## Acknowledgments

This work was supported by the Hartman Foundation for Parkinson’s disease.

## Notes

**Authors report no conflict of interest**

